# GO Bench: Shared-hub for Universal Benchmarking of Machine Learning-Based Protein Functional Annotations

**DOI:** 10.1101/2022.07.19.500685

**Authors:** Andrew Dickson, Ehsaneddin Asgari, Alice C. McHardy, Mohammad R.K. Mofrad

**Affiliations:** Molecular Cell Biomechanics Laboratory, Departments of Bioengineering and Mechanical Engineering, University of California, Berkeley, CA, 94720, USA; Computational Biology of Infection Research, Helmholtz Centre for Infection Research, Brunswick 38124, Germany

**Author notes:** Equal contribution (Andrew Dickson, Ehsaneddin Asgari).

## Abstract

**Motivation:** Gene annotation is the problem of mapping proteins to their functions represented as Gene Ontology terms, typically inferred based on the primary sequences. Gene annotation is a multi-label multi-class classification problem, which has generated growing interest for its uses in the characterization of millions of proteins with unknown functions. However, there is no standard GO dataset used for benchmarking the newly developed new machine learning models within the bioinformatics community. Thus, the significance of improvements for these models remains unclear.

**Summary:** The Gene Benchmarking database is the first effort to provide an easy-to-use and configurable hub for the learning and evaluation of gene annotation models. It provides easy access to pre-specified datasets and takes the non-trivial steps of preprocessing and filtering all data according to custom presets using a web interface. The GO bench web application can also be used to evaluate and display any trained model on leaderboards for annotation tasks.

**Availability and Implementation:** The GO Benchmarking dataset is freely available at llp.berkeley.edu/GO_bench/dataset_form, and code is available at http://github.com/amdson/GO_pipeline.

**Supplementary information:** Supplementary data are available at *Bioinformatics* online.

## 1 Introduction

The Gene Ontology (GO), is a curated network of over 40,000 ‘terms’ capturing possible functions of a protein Ashburner *et al.* (2000). Matching a protein with one or more GO terms is a useful method for characterization, but out of hundreds of millions of sequenced proteins, only a small fraction are functionally annotated. Thus, a goal for the bioinformatics community is predicting annotations from sequence, with potential to aid researchers in understanding organism proteomes (Ghatak *et al.*, 2019) or identifying the significance of mutations (Ding *et al.*, 2004). Dozens of ML methods have emerged for GO annotation (Zhou *et al.*, 2019), but there are several barriers to evaluating new models, starting with the technical difficulty of generating a strong dataset with relevant metrics. Creating a complete dataset often requires downloading and parsing the 90 GB Gene Ontology Annotation database and developing custom scripts. Additionally, it is difficult to immediately compare results from models built on different datasets. The current platform for fair comparison of GO annotation models is the CAFA competition (Zhou *et al.*, 2019), but because this relies on new experimental data for evaluation, it is only possible on a semi-annual basis and may have varying difficulty. Typically, current teams simply reproduce their efforts for each model they wish to compare against, or replicate the CAFA dataset. In both cases, this is a costly and inefficient process. Our GO Bench website offers several simple and standardized datasets for the GO annotation problem, and allows users to customize their datasets through a simple interface while still allowing for comparisons between results. It also allows researchers to upload model predictions, in the same style as for the CAFA competition, and view benchmarking statistics for all submitted models.

## 2 Features

GO Bench’s primary purpose is to generate customizable but compatible datasets of GO annotations, based on user specified pipeline settings. In addition, GO Bench allows users to upload model predictions in order to compare their results on a public leaderboard. **Data Collection:** 566,996 high-quality protein sequences are obtained from SwissProt (Consortium, 2020), while the Gene Ontology Annotation (GOA) dataset is cross-referenced for roughly 1.3 million annotations (Huntley *et al.*, 2015). Annotations are associated with evidence codes representing their quality (Chibucos *et al.*, 2017) and filtered for experimental evidence only by default. However, there is the option to include lower-quality annotations, such as those that are predicted, but human-reviewed, because they may improve model performance in practice. **Train-Test:** For model evaluation following training, we pre-generate two data divisions, with one split assigning proteins to training or testing randomly, and the other grouping proteins with high (≥ 50%) similarity. The latter is used by default since it prevents highly similar proteins from appearing in training and testing and artificially inflating results. For compatibility, proteins used in “Benchmarking gene ontology function predictions using negative annotations” are filtered from testing data(Warwick Vesztrocy and Dessimoz, 2020). **Annotation Propagation:** The gene ontology contains functional terms, but also an inheritance structure in which “child” terms qualify as instances of their “parents” (Ashburner *et al.*, 2000). Generally, only the most specific term is recorded when a protein annotation is discovered, thus, annotations in a dataset are optionally propagated up through the gene ontology structure before use. **Term Filtering:** The gene ontology contains over forty thousand terms, and the most specific terms have few or even zero associated proteins. GO Bench typically filters out terms which are too infrequent for a model to process based on a user determined cutoff. Other factors, such as the quality of evidence for an annotation or the date at which an annotation was generated, are also available for filtering. **Leaderboard:** Users may submit model predictions to a leaderboard containing several different testing datasets and metrics. In particular, our leaderboard evaluates models on the *S_min_* and *F_max_* metrics used by the popular CAFA competition (Zhou *et al.*, 2019;Clark and Radivojac, 2013), as well as on the F_score_ metric known to work well for datasets with many positive, but unlabeled datapoints (Bekker and Davis, 2020) (§S-5.3). Plots of model performance are displayed for each leaderboard entries.

## 3 Results and Conclusion

We include baseline models including the CAFA version of BLAST, a convolutional model based on DeepGOPlus (Kulmanov and Hoehndorf, 2019), and a feedforward network with 3-mer distributions of sequences as an input, similar to the original ProtVec paper (Asgari and Mofrad, 2015). The naive model is based solely on GO annotation frequencies. In table (a), we’ve displayed our results for the Molecular Function category with a cluster50 train/test split, and only experimental evidence codes in testing data. Notably, our baseline 3-mer model performs best when trained on all but completely computational annotations. To our knowledge, the GO Bench project provides the first directly usable datasets for training and benchmarking GO annotation models. They allow for flexibility in trade-offs, but maintain a backbone of the same testing data split to guarantees compatibility between results. Based on these, we offer competitive baselines, and find that previously untapped data sources can substantially improve performance. We expect GO Bench to be widely used by the bioinformatics community because it unifies and simplifies efforts to evaluate new models through a standard benchmark presented in the GO Bench leaderboard.

**Table (a).** Baseline Results for GO Annotation. Models were evaluated in the Molecular Function domain on the *F_max_*, *S_min_*, and *F_score_* metrics (detailed in §S.1.1), and train/test split is by cluster (§S.2). The 3-mer model was trained on experimental annotations only, any human-reviewed annotations (non-IEA), and finally all annotations. Highest scores are bolded for each metric.

| Model | Molecular Function |  |  |
| --- | --- | --- | --- |
| | $F_{max}$ | $S_{min}$ | $F_{score}$ |
| 3-mer (EXP) | 0.53 | 11.83 | 216.48 |
| 3-mer (non-IEA) | 0.59 | 10.73 | 255.09 |
| 3-mer (ALL) | 0.51 | 12.57 | 192.21 |
| **3-mer (BLAST)** | **0.6** | 10.58 | 263.78 |
| DPG Conv (non-IEA) | 0.6 | **10.5** | **267.67** |
| BLAST | 0.42 | 13.62 | 131.89 |
| Naive | 0.42 | 13.4 | 153.61 |

**Figure (b).**
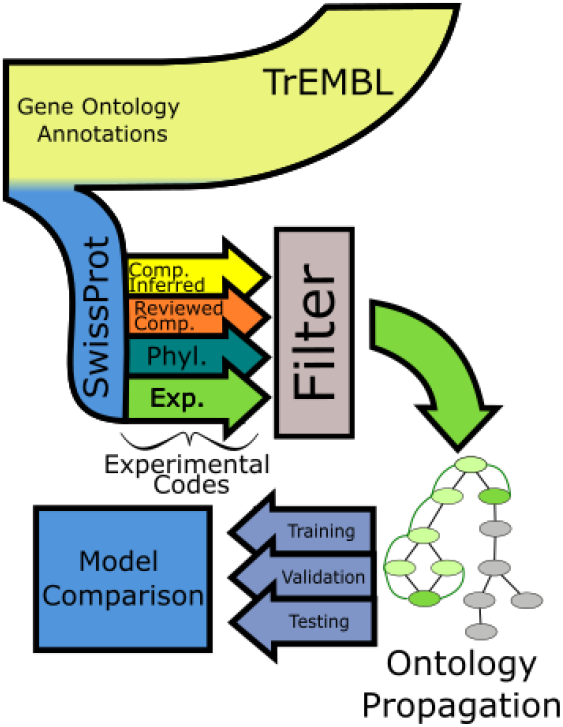
The GO Bench data pipeline extracts annotated SwissProt sequences from TrEMBL, and categorizes annotations by quality for user filtering. Annotations are propagated up the Gene Ontology tree structure and split into training, validation, and testing datasets.

## Supporting information

Supplementary Information for "GO Bench: Shared-hub for Universal Benchmarking of Machine Learning-Based Protein Functional Annotations"

## Acknowledgements

This work has been supported by the Open Computing Facility at UC Berkeley.

