## Supplementary Information for "GO Bench: Shared-hub for Universal Benchmarking of Machine Learning-Based Protein Functional Annotations" for "GO Bench: Shared-hub for Universal Benchmarking of Machine Learning-Based Protein Functional Annotations"

picture(0,0)(-35,0)(1,0)30 (0,35)(0,-1)30 picture picture(0,0)(35,0)(-1,0)30 (0,35)(0,-1)30 picture

*Bioinformatics*
doi.10.1093/bioinformatics/xxxxxx
Advance Access Publication Date: Day Month Year
Application Note

OXFORD

Sequence Analysis

### GO Bench: Shared-hub for Universal Benchmarking of Machine Learning-Based Protein Functional Annotations

**Andrew Dickson*[1], Ehsaneddin Asgari*[1,2], Alice C. McHardy[2], and Mohammad R.K. Mofrad[1†]**

[1]Molecular Cell Biomechanics Laboratory, Departments of Bioengineering and Mechanical Engineering, University of California, Berkeley, CA, 94720, USA
[2]Computational Biology of Infection Research, Helmholtz Centre for Infection Research, Brunswick 38124, Germany

* Equal contribution.
† To whom correspondence should be addressed.

Associate Editor: XXXXXXX

Received on XXXXX; revised on XXXXX; accepted on XXXXX

#### Abstract

**Contact:**
**Supplementary information:** Supplementary information contained here.

#### 1 Data Preparation

Annotation data and the gene ontology itself are downloaded from the Gene Ontology Association (GOA) database and processed with in-house scripts developed in Python. Code is included in https://github.com/amdson/go_bench. For model training, protein identifiers and sequences are downloaded separately from UniProt's SwissProt database.

The GOA database is the April 2021 version from "ftp://ftp.ebi.ac.uk". It takes the form of a tab separated CSV file, with fields for UniProt protein ID, Gene Ontology annotation, date of annotation, and annotation qualifiers such as "NOT", "contributes to", or "colocalizes with". Each row of the GOA database represents a single annotation for a specific protein.

The entire dataset is prefiltered to removed annotations not for Swissprot proteins. Any annotations with negative qualifiers are also removed. All GO Bench datasets are sourced from a local version of the filtered GOA dataset. For model training, and to filter non-Swissprot proteins, we use the August 2021 version of Uniprot, located at "ftp.uniprot.org/pub/databases/uniprot/previous_major_releases".

#### 2 Train-Test Split

We provide two train-validation-test splits for user downloads, named cluster50 and random.

All sequences used for our cluster50 datasets are extracted from the UniRef50 dataset (SwissProt), in which sequences are clustered to 50% identity. Clusters are randomly assigned to one of training, validation, or testing categories until they are as close as possible to fractions of 70, 15, and 15 percent respectively. This guarantees that proteins with over 50% similarity are never assigned to different portions of the dataset, which would compromise the integrity of several validation functions. Our random dataset is primarily for reference, and is generated using the same proteins, but without accounting for clustering. Splits are entirely random, but have the same proportions.

Once users select their train-test split, datasets may be further customized with operations such as annotation propagation, filtering, or the addition of estimated negative annotations. However, regardless of customizations, the underlying train-test split remains the same, so that the resulting models can be safely added to the corresponding leaderboard.

#### 3 Model Architecture

Benchmark models were developed using Pytorch, the BLAST package, and BLAST annotation scripts provided by CAFA. All code is included in the on github at github.com/amdson/GO_benchmarking.

BLAST annotations were generated using the same process and methodology as in CAFA, but with our own training data.

For our deep learning benchmarks, we chose several basic models, a feedforward neural network taking in a 3-mer embedding, an LSTM taking

© The Author 2018. Published by Oxford University Press. All rights reserved. For permissions, please 

picture(0,0)(-35,0)(1,0)30 (0,-35)(0,1)30 picture picture(0,0)(35,0)(-1,0)30 (0,-35)(0,1)30 picture
"output" — 2022/6/22 — page 1 — #1

in sequences directly, and a convolutional model also taking in sequences directly.

Our 3-mer embedding was generated by feeding our training dataset into the tf-idf transform from the sklearn python library. The transform extracts the count of every possible sequential triple of amino acids in a sequence, and then downweights by a heuristic measure to prevent the most common triples from dominating. This was then combined with the mean of ProtVec embeddings of all 3-mers in a protein to generate a composite vector of roughly 10,000 entries.

The feedforward network is implemented in Pytorch. It contains one input layer, two hidden layers of size 5000, and one output layer generating probabilities for each GO term in parallel. Between each layer, we used Dropout with rate 0.5, BatchNorm regularizers and ReLU non-linearities. For each GO domain, we use the same network architecture, and a network is trained for each domain.

Our convolutional network is built implemented in Pytorch, using methodology from the network component of the Deep Pred GO model. It is a shallow convolutional model which applies 1D convolutional filters ranging in size from 3 to 80 the entire protein sequence, and then applies a fully-connected feedforward layer to their concatenated output. Full details of the convolutional implementation are included in the DeepGoPlus paper, as well as our codebase.

#### 4 Model Training

Deep learning models are trained using the Pytorch Lightning package. Each uses Lightning's heuristically determined learning rate on a linearly decaying schedule, and the Adam optimizer for gradient descent. For all networks, we use a batch size of 128 All networks are trained on a GeForce 1080 GPU until validation performance decreases for three successive epochs.

#### 5 Performance Metrics

GO Bench uses the F-max, S-min, and F-score metrics mentioned previously. The F-max and S-min metrics are directly based off of the CAFA competition, and are described in detail in the CAFA 2 paper "An expanded evaluation of protein function prediction methods shows an improvement in accuracy" (Jiang and Oron, 2016). The 'F-Score' metric is defined in "Learning from Positive and Unlabeled Data: a survey" (Bekker and Davis, 2020). For clarification, we describe all metrics below.

Let $G_n$ represent the set of GO terms in a given namespace $n$, and $P$ represent the set of proteins within our dataset. For the purpose of metric evaluation, we write GO annotation as a binary classification problem, in which a pair $x = (g \in G_n, p \in P)$ is classified as either true or false.

Define classification models as functions parameterized by $\theta$ mapping GO term, protein pairs to probability estimates. $M_\theta | G_n \times P \to [0, 1]$. At a given threshold $\alpha$, the classification estimate for an input $x = (g, p)$ is $\hat{y} = M_\theta(x) \geq \alpha$.

GO Bench datasets consist of positive GO annotation for a a given set of proteins (within the constraints of customization). Negative annotation is represented implicitly by absence from the dataset. We write the complete dataset as $D = (X, y)$, where $X$ represents all pairs of proteins included in the dataset and GO terms, and $y \in \{0, 1\}$ gives the classification label for that pair. Assume there are $N$ datapoints, with $(x_i, y_i)$ giving the ith pair.

##### 5.1 F-Max

For a given $\alpha$, define precision as the number of true positive predictions divided by the number of positive predictions and recall as the number of true positive predictions divided by the number positive annotations.

$$pr(\alpha) = \frac{\sum_{i=1}^{N} 1(y_i = \hat{y}_i(\alpha))}{\sum_{i=1}^{N} \hat{y}_i(\alpha)} \quad (1) \qquad rc(\alpha) = \frac{\sum_{i=1}^{N} 1(y_i = \hat{y}_i(\alpha))}{\sum_{i=1}^{N} y_i} \quad (2)$$

The F1 score is defined as $\frac{2pr(\alpha)rc(\alpha)}{pr(\alpha)+rc(\alpha)}$, and the F-max score is $\max_{\alpha \in [0,1]} F1(\alpha)$. When calculating the F1 score, we use the 'micro-F1' score defined in the python sklearn package (Pedregosa *et al.*, 2011), and evaluate results over threshold values ranging from 0 to 1 over increments of 0.01 to take the maximum.

##### 5.2 S-min

Information content is a quantitative measure of the descriptive value of a gene ontology term, defined using the GO graph structure (Clark and Radivojac, 2013). It associates a value $ic(g)$ to each GO term $g \in G_n$.

With information content, define remaining uncertainty and missing information as the following.

$$ru(\alpha) = \frac{1}{|P|} \sum_{i=1}^{N} ic(g_i) \cdot 1(\hat{y}_i(\alpha) = 0 \wedge y_i = 1) \quad (3)$$

$$mi(\alpha) = \frac{1}{|P|} \sum_{i=1}^{N} ic(g_i) \cdot 1(\hat{y}_i(\alpha) = 1 \wedge y_i = 0) \quad (4)$$

The final S-min score is $S_{min} = min_\alpha \left( \sqrt{ru(\alpha)^2 + mi(\alpha)^2} \right)$.

##### 5.3 F-Score

The F-Score metric is defined to approximate the true F1 score with perfect knowledge of datapoint labels (Bekker and Davis, 2020). It's based on recall, $rc(\alpha)$, and the probability of a positive prediction, $P(\hat{y}(\alpha) = 1) = \frac{1}{N} \sum_{i=1}^{N} \hat{y}(\alpha)$. For a given $\alpha$, the F-Score is

$$\frac{rc(\alpha)^2}{P(\hat{y}(\alpha) = 1)}$$

As with F-max, we take the maximum over all thresholds $\alpha$ for our results.

#### 6 Website

The GO Bench website is hosted at llp.berkeley.edu/GO_bench/dataset_form. It contains a simple webform for customizing and downloading datasets, visualizations of class frequencies within downloaded datasets, a leaderboard for submitted models, and scoring visualizations for uploaded models.

picture(0,0)(-35,0)(1,0)30 (0,35)(0,-1)30 picture
picture(0,0)(35,0)(-1,0)30 (0,35)(0,-1)30 picture



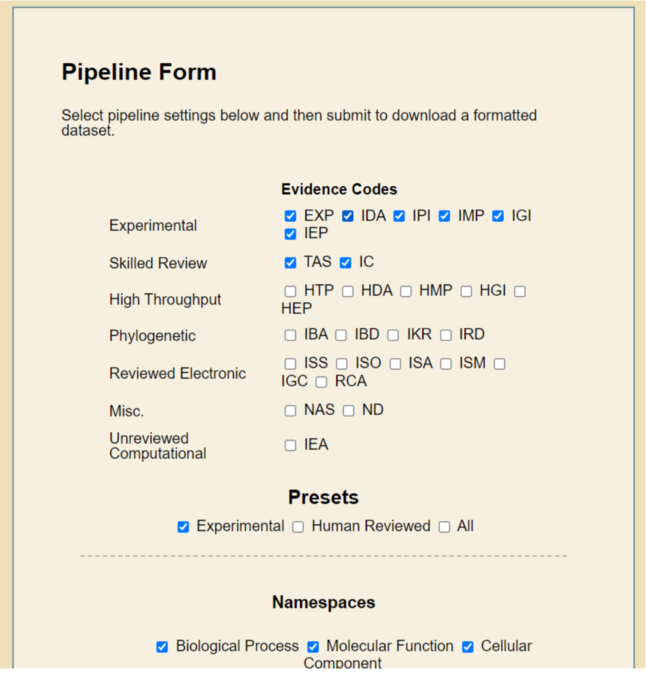


Fig. 1: Initial form for generating images. Form is built from the HTML <form> element, and contains entries for annotation quality, namespaces, annotation propagation, term filtration, train-test split method, and cutoff dates for annotation assignment.

picture(0,0)(-35,0)(1,0)30 (0,-35)(0,1)30 picture
picture(0,0)(35,0)(-1,0)30 (0,-35)(0,1)30 picture
“output” — 2022/6/22 — page 3 — #3

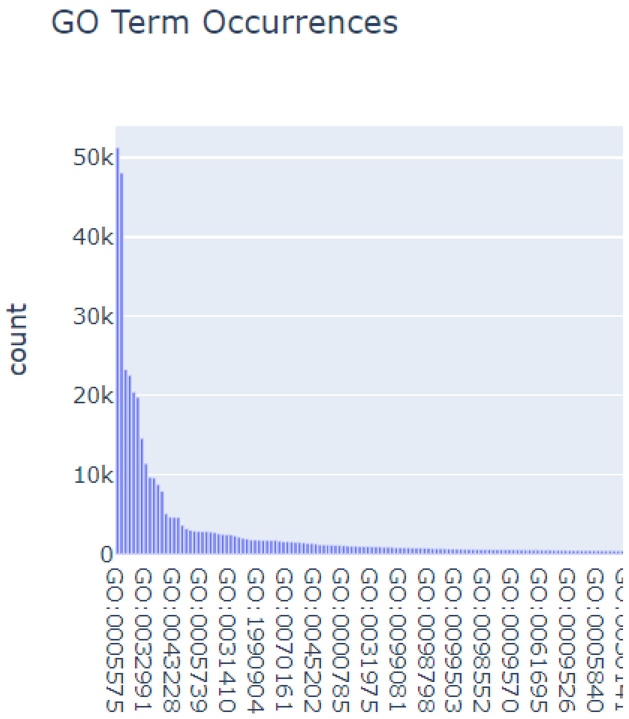


Fig. 2: Histogram visualization of GO class counts for each GO entry. Separate counts are given for each namespace in the gene ontology (Molecular Function, Cellular Component, and Biological Process). Common terms such as protein interaction appear in tens of thousands of dataset proteins, while the rarest ones appear only once, and are usually filtered out by form settings.

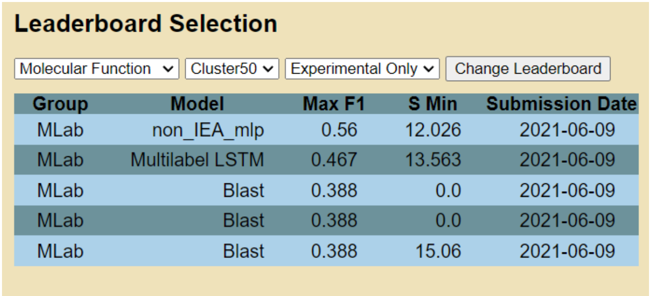
**Leaderboard Selection**

Molecular Function | Cluster50 | Experimental Only | Change Leaderboard

| Group | Model | Max F1 | S Min | Submission Date |
|---|---|---|---|---|
| MLab | non_IEA_mlp | 0.56 | 12.026 | 2021-06-09 |
| MLab | Multilabel LSTM | 0.467 | 13.563 | 2021-06-09 |
| MLab | Blast | 0.388 | 0.0 | 2021-06-09 |
| MLab | Blast | 0.388 | 0.0 | 2021-06-09 |
| MLab | Blast | 0.388 | 15.06 | 2021-06-09 |

Fig. 3: The GO Bench leaderboard records F-Max, S-Min, and F-Score metrics for each uploaded model, and presents a top ten list for the highest ranking versions.

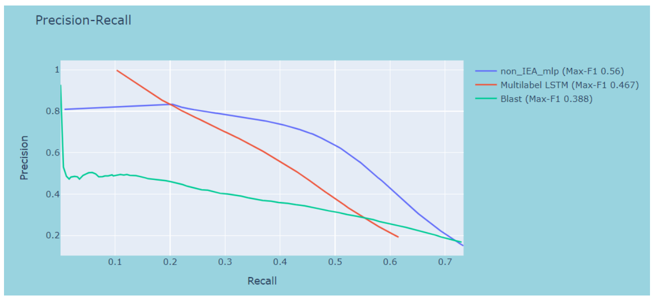


Fig. 4: The GO Bench leaderboard is also be visualized using plots of scoring metrics including precision-recall.
